# Comprehensive, high-resolution binding energy landscapes reveal context dependencies of transcription factor binding

**DOI:** 10.1101/193904

**Authors:** Daniel D. Le, Tyler C. Shimko, Arjun K. Aditham, Allison M. Keys, Yaron Orenstein, Polly M. Fordyce

## Abstract

Transcription factors (TFs) are primary regulators of gene expression in cells, where they bind specific genomic target sites to control transcription. Quantitative measurements of TF-DNA binding energies can improve the accuracy of predictions of TF occupancy and downstream gene expression *in vivo* and further shed light on how transcriptional networks are rewired throughout evolution. Here, we present a novel sequencing-based TF binding assay and analysis pipeline capable of providing quantitative estimates of binding energies for more than one million DNA sequences in parallel at high energetic resolution. Using this platform, we measured the binding energies associated with all possible combinations of 10 nucleotides flanking the known consensus DNA target for two model yeast TFs, Pho4 and Cbf1. A large fraction of these flanking mutations change overall binding energies by an amount equal to or greater than consensus site mutations, suggesting that current definitions of TF binding sites may be too restrictive. By systematically comparing estimates of binding energies output by deep neural networks (NN) and biophysical models trained on these data, we establish that dinucleotide specificities are sufficient to explain essentially all variance in observed binding behavior, with Cbf1 binding exhibiting significantly more epistasis than Pho4. NN-derived binding energies agree with orthogonal biochemical measurements and reveal that dynamically occupied sites *in vivo* are both energetically and mutationally distant from the highest-affinity sites.

---

Gene expression is extensively regulated by transcription factors (TFs) that bind genomic sequences to activate or repress transcription of target genes (1). The strength of binding between a TF and a given DNA sequence at equilibrium depends on the change in Gibbs free energy (ΔG) of the interaction (2–5). Thermodynamic models that explicitly incorporate quantitative estimates of binding energies more accurately predict occupancies, rates of transcription, and levels of gene expression *in vivo* (4, 6–11). In addition, binding energy measurements for TF-DNA interactions can provide insights into the evolution of regulatory networks. Unlike coding sequence variants that manifest at the protein level to influence fitness, non-coding TF target site variants affect phenotype by modulating the binding energies of these interactions to affect gene expression (12, 13). Understanding how TFs identify their cognate DNA target sites *in vivo* and how these interactions change during evolution therefore requires the ability to accurately estimate binding energies for a wide ranges of sequences.

Most high-throughput efforts to develop accurate models of TF binding specificities have focused on the effects of mutations within known TF target sites that dramatically change binding energies. However, even subtle changes in binding energies can have dramatic effects on both occupancy and levels of transcription (14–17). Sequences surprisingly distal from a known consensus motif can affect affinities and levels of transcription (18–21), and genomic variants in regulatory regions outside of known TFBSs may be subject to non-neutral evolutionary pressures (22). Therefore, understanding the fundamental mechanisms that regulate transcription requires the ability to not only measure binding energies for a large number of sequences, but to do so at sufficient resolution to resolve even small effects.

Despite the utility of comprehensive binding energy measurements, existing characterization methods often lack the energetic resolution and scale required to yield such datasets. Currently, three dominant technologies are used to query DNA specificities: methods based on systematic evolution of ligands by exponential enrichment (SELEX) (23–28), protein binding microarrays (PBMs) (29, 30), and mechanically-induced trapping of molecular interactions (MITOMI) (31, 32). SELEX-based methods require repeated enrichment and amplification cycles followed by sequencing of the enriched TF-bound material. Thus, these methods are optimized to identify the highest affinity substrates from extremely large random populations, but fail to measure moderately- or weakly- bound sequences, much less measure their affinities. Although PBMs quantify the binding of TFs to DNA microarrays using a fluorometric readout with a broad dynamic range, the precise relationship between measured fluorescence intensities and binding energies is unclear because PBMs require wash steps that disrupt binding equilibrium. The MITOMI platform, based on mechanical trapping of molecular interactions by microfluidic valves, enables high-resolution measurements of concentration-dependent to yield absolute affinities, but is limited to characterization of several hundred sequences. Recent iterations of MITOMI-based binding assays have addressed these throughput limitations by employing massively parallel sequencing to increase sequence space coverage, but at the cost of resolving binding energies (28, 33). Other techniques such as high-throughput sequencing-fluorescent ligand interaction profiling (HiTS-FLIP) couple massively parallel sequencing with the ability to perform concentration-dependent binding measurements; however, adoption of this technology has been limited by the requirement for extensively customized sequencing hardware (34). Taken together, these TF-DNA binding assays can sample vast sequence spaces, but it remains challenging to simultaneously measure binding energies at the scale and resolution necessary to derive complete binding energy landscapes.

## Significance Statement

Transcription factors (TFs) are key proteins that bind DNA targets to coordinate gene expression in cells. Understanding how TFs recognize their DNA targets is essential for understanding how variations in regulatory sequence disrupt transcription to cause disease. Here, we develop a novel high-throughput assay and analysis pipeline capable of measuring absolute binding energies for over one million sequences with high resolution, and apply it towards understanding how nucleotides flanking DNA targets affect binding energies for 2 model yeast TFs. Through systematic comparisons between models trained on these data, we establish that considering dinucleotide interactions is sufficient to accurately predict binding, and further show that sites used by TFs *in vivo* are both energetically and mutationally distant from the highest-affinity sequence.

The energetic resolution of direct binding energy measurements is fundamentally limited to ∼1 RT (or ∼0.5 kcal/mol) by intrinsic thermal noise. However, measuring binding to a large library of sequences can yield even higher precision estimates of binding energies for individual sequences by training specificity models on large amounts of noisy data. The most popular and widely used models represent TF specificities as a position weight matrix (PWM), in which each nucleotide at each position contributes additively and independently to overall binding energies (35). These mononucleotide models are easily implemented, visualized, and interpreted, and provide useful approximations of binding specificity for the majority of studied TFs (9, 36–38). However, PWM-based models fail to capture epistasis between nucleotides, which can lead to inaccurate predictions, particularly for low-affinity sites (31, 39). This approximation can be further refined by including contributions of higher-order sequence features, such as dinucleotides or longer k-mers (40–49). Several recently developed mechanistic models predict binding based on biophysical properties of DNA, such as the local shape of DNA sequences (*e.g.* minor groove width, propeller twist, helical twist, and roll) (50–54). However, these biophysical variables are predicted directly from primary sequence, rendering the relationship between the two somewhat degenerate. Deep neural network (NN)-based models have shown considerable success in learning complex patterns from large datasets across a variety of applications, including predicting the function of non-coding genomic sequences (55). Training NN models on large sets of binding data therefore has the potential to yield more accurate and higher-resolution estimation of binding behaviors at a per-sequence level, revealing local topography on of binding energy landscapes.

To address the need for technologies capable of highthroughput thermodynamic measurements, we developed an integrated high-throughput sequencing assay and analysis pipeline capable of estimating both relative and absolute binding energies (ΔΔG and ΔG, respectively) for >1 million sequences in parallel, even for relatively small energetic differences. Using Monte Carlo simulations designed to mimic the effects of stochastic sampling noise on energetic resolution, we establish guidelines for the sequencing depth required to resolve accurate binding energies for libraries of different sizes and expected energy ranges. We then deploy this assay to measure comprehensive and quantitative binding energy landscapes for >1 million mutations surrounding the known consensus motif for two model yeast TFs (Pho4 and Cbf1). Deep neural network (NN) models that incorporate all possible higher-order, non-additive contributions were then trained on these large datasets to yield high-resolution estimates of binding specificity for each sequence. Comparisons to orthogonal biochemical affinity measurements established that NN predictions are highly quantitative, accurately predicting measured binding energies over a range of 3 kcal/mol. A surprisingly large number of sequences tested have effects on binding energies as great or greater than mutations in the core, suggesting that current definitions of TFBSs are too restrictive and may limit accurate predictions of TF occupancy *in vivo*. Comparisons between NN-derived predictions and predictions derived from a series of biophysically motivated models reveal that dinucleotide specificity preferences are sufficient to explain nearly all observed binding behavior, with Cbf1 exhibiting significantly more epistasis than Pho4. Strikingly, most dynamically occupied target loci for both Pho4 or Cbf1 are mutationally distal from the energetically optimal flanking sequence representing the global minimum of the energy landscape, providing evidence of evolutionary molecular selection for near-neutral effects on binding energies. Taken together, these data demonstrate the utility of our high-throughput approach to measure relative binding energies and model determinants of substrate specificity required to understand biological behaviors. Furthermore, the experimental design and analysis frameworks presented here may be extended to a wide variety of currently incompletely characterized TFs, improving predictive models of TF-DNA affinities across species.

## Results

### A microfluidic approach employing high-throughput sequencing to derive comprehensive binding affinity landscapes

We sought to develop an assay that significantly extends the scale at which TF-DNA interactions can be probed while maintaining the ability to quantitatively measure binding energies at high resolution. TF-DNA interactions can be considered a two-state system, such that the affinity of a given interaction can be determined by the equilibrium partitioning of sequences into bound and unbound states:

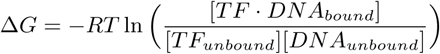

In recent work, several groups have established that molecular counting of individual DNA molecules via high-throughput sequencing can reliably measure bound and input concentrations for each species (10, 39, 56). These assays generally employ electromobility shift assays (EMSAs) to isolate bound material. However, TF-DNA complexes are not at chemical equilibrium during the electrophoresis step and complexes with particularly fast dissociation rates may be underrepresented within the bound fraction, leading to a systematic underestimation of weak affinity interactions (57). To address this issue, we employed a microfluidic device incorporating pneumatic valves with fast (∼100 ms) actuation times to mechanically “trap” DNA associated with TF proteins at equilibrium (28, 31– 33) (Fig. 1*A*). This device requires very small amounts of both DNA substrate and expressed protein, eliminating the need for cell-based protein production. Antibody-patterned surfaces within the device capture meGFP-tagged TFs produced via *in vitro* transcription/translation prior to washing, effectively purifying the protein *in situ*. After TF capture, libraries of DNA sequences are introduced and allowed to interact with surface-immobilized TFs until equilibrium is reached. Mechanical valves then sequester TF-bound DNA sequences, making it possible to wash out unbound material without loss of weak interactions (31, 32, 58) (Fig. 1*B*). Isolated bound DNA species can then be eluted from the device and quantified using highthroughput sequencing (Fig. 1*C*). The concentration of DNA within the device (∼1 μM) is in excess of the immobilized TF concentration (∼30 nM), allowing approximation of unbound concentrations via sequencing of the input library.

**Fig. 1.**
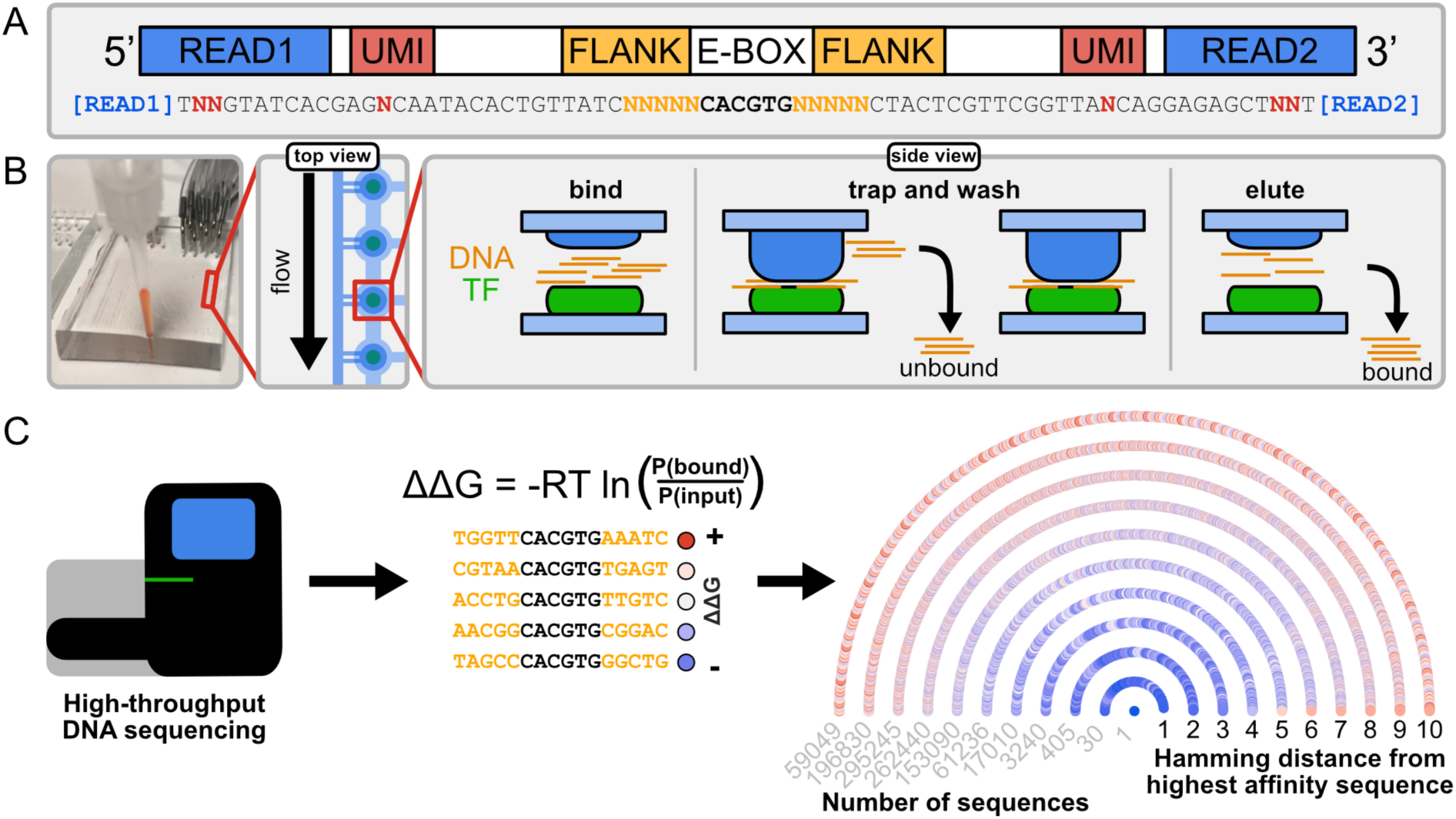
DNA library design and assay overview. (*A*) Schematic of flanking sequence library design indicating locations of Illumina sequencing adapters (blue), unique molecular identifiers (UMIs, red), variable flanking regions (orange), and E-box consensus. (*B*) Photograph of MITOMI device (left) and schematic showing device operation (right). (*C*) Schematic showing downstream sample analysis. Counting of individual molecules within bound and input fractions allows calculation of relative binding energies for each sequence (left, middle) to yield a comprehensive thermodynamic binding affinity landscape (Pho4 example). Each color-coded point (blue = −ΔΔG, red = +ΔΔG) represents a sequence, grouped by Hamming distance from the highest affinity sequence and alphabetically ordered in clockwise polar coordinates.

As a first application of this assay, we focused on two model TFs from *Saccharomyces cerevisiae*, Pho4 and Cbf1. Although Pho4 and Cbf1 are known to bind the same CACGTG variant of the 6-nucleotide enhancer-box (E-box) motif both *in vitro* and *in vivo* (15, 17, 27, 59–61), they bind largely nonoverlapping sets of genomic loci and regulate distinct sets of target genes (18, 62). To comprehensively probe how flanking nucleotides affect Pho4 and Cbf1 binding affinities, we designed a library of 1,048,576 sequences in which the core E-box sequence was flanked by all possible random combinations of five nucleotides upstream and downstream, embedded within a constant sequence empirically shown to exhibit negligible binding (32) (Fig. 1*A*). Because only sparse quantities of material are eluted from the device at the end of each experiment, eluted material must be amplified by PCR prior to Illumina sequencing. Constant sites at the 5’ and 3’ ends allowed simultaneous PCR amplification and incorporation of Illumina adapters, and unique molecular identifiers (UMIs) included within each library sequence allowed accurate counting of library species even in the presence of PCR bias (63). Each UMI barcode was segmented and interspersed along the library sequence to prevent the formation of an additional CACGTG consensus site. After sequencing, relative binding affinities (ΔΔGs) were calculated for all sequences by considering relative enrichment of individual DNA species in the TF-bound fraction compared to the input library, thereby generating a comprehensive binding affinity landscape (example shown in Fig. 1*C*).

### Assay simulations determine sequencing depth requirements for binding affinity measurements

Accurately estimating concentrations of DNA in TF-bound and input samples via sequencing requires that measured read counts reflect true abundances. However, read counts can be distorted by stochastic sampling error, particularly for low read count numbers (64, 65). To understand how stochastic sampling error depends on read depth, library size, and the expected range of binding energies across library sequences, we began by considering a previously published experiment that quantified interactions between the *E. coli* LacI repressor and a library of 1,024 binding site variants via deep sequencing (library R3.2) (39). Each sequence was sampled to an average depth of roughly 10^3^ reads per species, yielding ΔΔG measurements with negligible sampling noise. To understand how read depth affects the recovery of accurate ΔΔG measurements, we downsampled these data to simulate lower sequencing depths of 10^2^–10^6^ reads, split evenly between bound and input fractions (*ca.* 0.05–5,000 reads per sequence). We then assessed the accuracy of recovered ΔΔG values at these lower sequencing depths by calculating the squared Pearson’s correlation coefficient (*r*^2^) between ΔΔG values for each species calculated from downsampled data and published values calculated from the full data set. To minimize the effect of a few high accuracy values dominating the correlation statistic, each *r*^2^ was normalized by the fraction of observed species (Fig. 2*A*). For this 1,024-species library with binding energies that span ∼3 kcal/mol, ∼2 × 10^5^ total reads (∼100 reads per sequence) were sufficient to recover highly-accurate ΔΔG values for every sequence.

**Fig. 2.**
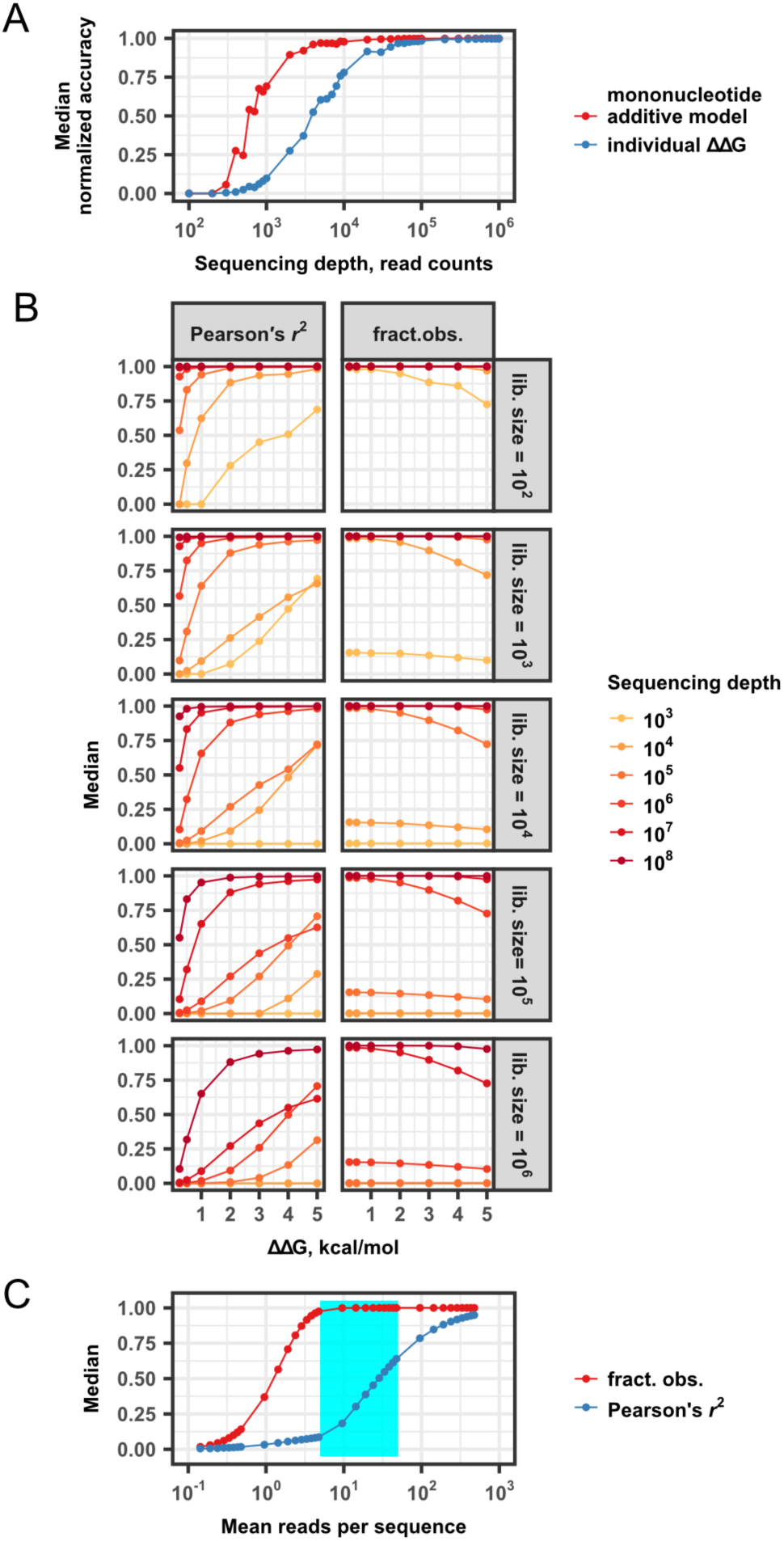
Probing the relationship between assay accuracy, read depth, library size, and energy range. (*A*) Normalized median accuracy (Pearson’s *r*^2^ * *f*, where *f* = the fraction of observed species) scores comparing results for downsampled data with “true” values as a function of read depth. The mononucleotide model reference values were based on PWM predictions that assume perfect additivity (39). Individual ΔΔG recovery (blue) and model recovery (red) shown. (*B*) Median squared Pearson’s correlation coefficients between recovered and “true” values (*r*^2^, left) and median fraction of observed species (right) as a function of binding affinity range for various library sizes (rows) sequenced to different depths. (*C*) Simulations parameterized to reflect putative flank library conditions. Median squared Pearson’s correlation coefficient (*r*^2^) between recovered and true values (blue) and fraction of observed species (red) as a function of mean reads per sequence for ten replicate simulations; a cyan rectangle denotes the assay configurations presented here.

Measuring accurate ΔΔGs for a 1,048,576 species library represents a 1000-fold increase in scale from these prior experiments. To understand more generally the determinants of ΔΔG measurement accuracy, we generated a simulated test set of “true” relative binding energies and implemented Monte Carlo simulations designed to mimic stochastic sampling during high-throughput sequencing. For given combinations of library sizes, sequencing depths, and binding energy ranges, we calculated the median squared Pearson’s correlation coefficient between calculated ΔΔG values and “true” values, as before. As expected, accuracy improves and library coverage expands with increasing sequencing depth (Fig. 2*B*, S1*A*, S1*B*). As the difference in expected binding energies increases, accuracy improves but the fraction of sequences observed from the input library decreases. Nearly all existing motif discovery libraries used in SELEX-type experiments probe on the order of 10^18^–10^24^ species with read depths of several thousand total reads. Extrapolating from the conditions considered in these simulations, such sparse sequencing depth relative to library size samples only the highest affinity sequences, representing an infinitesimal fraction of the input library. This limited set of observations likely misses smaller peaks within a binding energy landscape.

Previous observations of concentration-dependent Pho4 and Cbf1 binding to E-box motifs with mutations in the first flanking nucleotide revealed differences in affinities spanning ∼1 kcal/mol (31). To guide sequencing assays, we therefore examined in detail simulations sampling a 1,048,576-member library with this energy range at mean read depths per species ranging between 10^−1^–10^3^ (10^3^–10^8^ total reads) (Fig. 2*C*). Although 95% of sequences can be recovered from as few as 4–5 reads per species, high read depths of ∼10^2^ counts per species (10^8^ total reads per TF) are required to yield individual ΔΔG measurements with accuracies of −80% and errors of −0.2 kcal/mol.

### Modeling specificity from noisy individual measurements improves assay resolution

The requirement for high-depth sequencing may be cost-prohibitive for studies involving many TFs or when considering large DNA libraries. In those scenarios, modeling can be used to infer the determinants of binding specificity while minimizing stochastic sampling noise from sparse sequencing measurements. To illustrate the power of this approach, we again considered the published LacI repressor dataset (39). Although ∼10^2^ reads per sequence were required for accurate ΔΔG estimates, 10^1^ to 10^2^-fold fewer reads per sequence allowed the generation of additive mononucleotide PWM models with similar predictive power to those generated from the entire dataset (Fig. 2*A*). However, while PWMs yield reasonably good approximations of high affinity sequences, these models fail to explain the variance among lower affinity target sites that exhibit high sequence diversity (31, 39).

A neural network (NN) trained on millions of noisy individual per-sequence measurements can capture all measurable higher-order complexity, thereby yielding a high-resolution model capable of accurately predicting binding over a wide range of energies. However, this increased predictive power comes at the cost of interpretability. To improve the accuracy of our energetic estimates while preserving the ability to gain mechanistic insights, we applied a novel integrated measurement and modeling approach (Fig. 3*A*). First, we collected millions sequencing-based estimates of per-sequence ΔΔGs. Next, we trained a NN model on these sequencing data to obtain high-resolution energetic predictions for each substrate that capture the effects of all possible higher-order epistatic interactions among nucleotides. Finally, we parsed and quantified the biophysical mechanisms responsible for observed TF-DNA binding behaviors by systematically comparing correlations between predictions made by the NN model and a series of biophysically-motivated linear models (mononucleotide, nearest-neighbor dinucleotide, and all dinucleotide models). This integrated scheme simultaneously yielded a novel binding energy landscape of unprecedented scale and energetic resolution and allowed dissection of the biophysical mechanisms responsible for Pho4 and Cbf1 specificity.

**Fig. 3.**
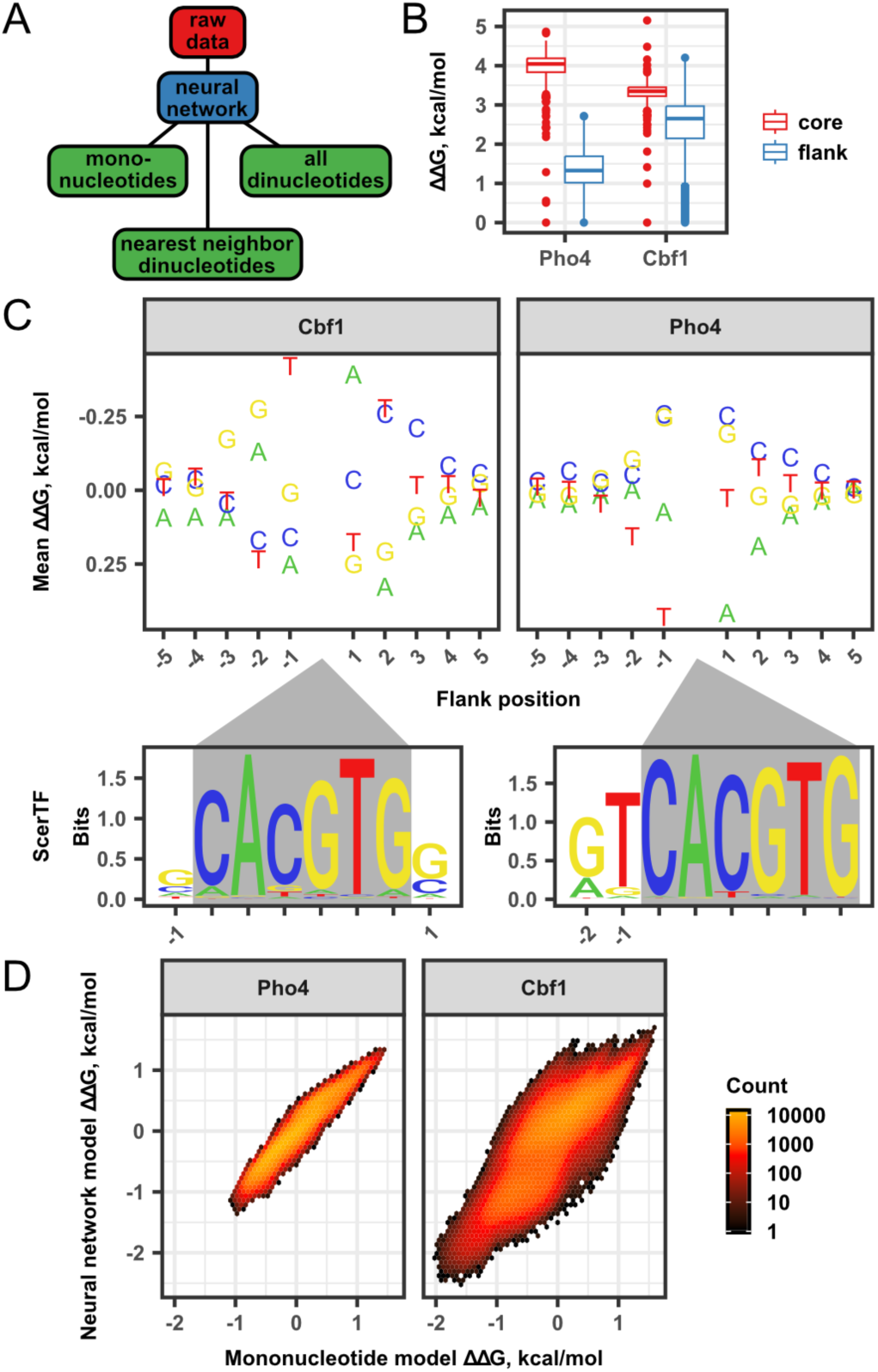
Modeling and interpretation of binding specificity based on mononucleotide features. (*A*) Data analysis flowchart shows neural network modeling from raw data, followed by interpretation based on linear combinations of sequence features. (*B*) Comparison of Pho4 and Cbf1 flanking sequence ΔΔG measurements with those derived from core mutations (31). (*C*) DNA base letters represent measured Pho4 and Cbf1 mean mononucleotide ΔΔ G measurements as a function of flanking sequence position, compared to previously derived sequence logos from the ScerTF database (66–69). Gray triangles and gray boxes represent position of the consensus E-box motif CACGTG. (*D*) For Pho4 and Cbf1, density scatter plots show the correlation between NN model estimates and mononucleotide additive model predictions based on those estimates.

### High-throughput, comprehensive estimates of absolute binding affinities for Pho4 and Cbf1

We used the assay and DNA library described above to acquire four replicate measurements of Pho4 and three of Cbf1 at sequencing depths ranging from approximately 5 to 50 million reads allocated to either bound or input samples (Tab. S1). For each experiment, ΔΔGs were calculated for each sequence from the measured ratio of bound to input read counts (Fig. S2). As predicted, measured per-sequence ΔΔGs between two experiments at low read depth (*ca.* 6–8 million limiting counts) show no correlation; conversely, at higher read depths (-24 million limiting counts), this correlation increases to *r*^2^ = 0.67 (Fig. S3, Tab. S2). To further improve resolution, we trained a NN regression model that predicted the measured ΔΔG for each input flanking sequence. Accuracy of the network against both the observed training data and an unobserved validation dataset was recorded throughout training. Training of the network was stopped once accuracy against the validation dataset failed to improve, protecting against overfitting to the training data (Fig. S4). Predictions from NN models trained on the 2 high-depth Pho4 replicates showed excellent correlation (*r*^2^= 0.94), validating the ability to apply such models to derive accurate, reproducible estimates of binding energies. For all analyses moving forward, we therefore use per-sequence ΔΔG estimates output from the NN trained on a composite dataset of all replicates (Fig. S5).

Estimates of absolute binding energy and dissociation constants (ΔG and K_d_, respectively) allow direct comparison between different TFs and across experimental platforms, and further enable quantitative predictions of TF occupancy *in vivo* under known cellular conditions. However, sequencing-based measurements of ΔGs from sparse datasets can underestimate the true affinity range due to systematic undersampling of bound reads for low-affinity sequences. In addition, given that the NN is trained only on relative binding affinities (ΔΔGs), it cannot return estimates of absolute energies (ΔGs). NNderived ΔΔG estimates can be projected onto an absolute scale by calibrating to a set of high-resolution biochemical measurements of ΔGs with a linear scaling factor and offset. To generate a set of high-confidence ΔGs, we applied the traditional fluorometric MITOMI assay to measure concentrationdependent binding behavior for surface-immobilized Pho4 and Cbf1 TFs interacting with all single-nucleotide variants of AGACA_TCGAG, a medium affinity reference flanking sequence (where the underscore indicates the CACGTG core motif) (Fig. S6*A*, S6*B*). For each oligonucleotide, observed binding was fit to a single-site binding model, yielding both K_d_s and ΔGs for each sequence (Tab. S3). The entire set of NN values was then scaled by fit parameters returned from a linear regression between NN predictions and experimental measurements for this set sequences. The median equilibrium dissociation constants for all flanking library sequences were 100 and 63 nM for Pho4 and Cbf1, respectively, in agreement with prior work (31). Strikingly, sequence variation in the flanking region around the consensus motif of Pho4 and Cbf1 can modulate K_d_ values by over two orders of magnitude, ranging between 11–1036 nM and 1–866 nM, respectively. In some cases, the magnitude of these effects exceeds that of mutations within the CACGTG core consensus (Fig. 3*B*, S7), demonstrating the importance of flanking sequences to specificity.

### Mononucleotide models reveal that Pho4 and Cbf1 flanking preferences extend far beyond the known consensus sequence

To understand the biophysical features that contribute to the predictive performance of the NN model, we generated PWMs (70), which estimate the mean energetic contribution of each nucleotide at each position, from the full set of scaled NN-predicted ΔΔG values (Fig. 3*C*). While the assumption of additivity fails to explain all specificity, these models often offer a close approximation (11, 43) and PWMs are easily visualized and interpreted. These mononucleotide model results confirm that positions proximal to the E-box core motif exhibit the largest mean effect on binding affinity, in agreement with PWMs generated by orthogonal techniques (66–68) (Fig. 3*C*). However, these results indicate that nucleotides up to 4 and 5 positions from the consensus contribute to specificity for Pho4 and Cbf1, respectively, significantly farther than previously reported.

To quantitatively assess the degree to which mononucleotide features dictate binding behavior, we determined the proportion of NN-derived per-sequence ΔΔG variance that is explained by a simple PWM (Fig. 3*D*). If mononucleotide models capture all determinants of observed specificity, PWM predictions would explain all the variance in NN-derived ΔΔG values. Conversely, discrepancies may indicate the presence of higher-order epistatic interactions not captured by the PWM. PWMs were capable of explaining a majority of the variance in NN predictions (*r*^2^ = 0.92 and *r*^2^ = 0.70 for Pho4 and Cbf1, respectively)(Tab. S4), consistent with the sentiment in the field that PWMs provide good approximations of sequence specificity (9, 11). Intriguingly, PWMs explain a significantly smaller proportion of observed Cbf1 measurement variance, suggesting that Cbf1 recognition may rely on higher-order determinants of specificity.

To evaluate assay reproducibility, we generated individual PWMs from each of the four Pho4 technical replicates and three Cbf1 technical replicates [Fig. S8]. Linear model coefficients for mononucleotides at each position were strongly correlated between replicates of a single TF (Pho4 *r*^2^ = 0.95– 0.97 and Cbf1 *r*^2^ = 0.79–0.87) and uncorrelated between TFs (Tab. S5, Fig. S9); a meGFP negative control protein exhibited no sequence specificity (Fig. S10). Using the fraction of unexplained variance (1-*r*^2^) as a precision metric, the expected error range in NN-derived mean mononucleotide ΔΔG values for Pho4 is 0.02–0.04 kcal/mol and that of Cbf1 is 0.09–0.16 kcal/mol. This analysis highlights the robustness of binding specificity models derived from the assay and data presented here.

### Dinucleotide models reveal that flanking nucleotides exhibit significant epistasis for Cbf1

The remaining unexplained variance observed between NN-derived values and PWM-predicted values (∼8% and ∼30% for Pho4 and Cbf1, respectively) could indicate the presence of higher-order epistatic interactions governing specificity, or could simply represent experimental noise (40) (Tab. S4). To probe for higher-order interactions, we fit two different dinucleotide models to the NN-derived scaled ΔΔG values: a nearest-neighbor model that explicitly considers contributions from adjacent dinucleotides and a more complex model that considers contributions from all dinucleotide combinations, including non-adjacent pairs (45). Direct comparisons between nearest-neighbor dinucleotide model-predicted values and NN-derived binding energies showed increased correlation for both Pho4 and Cbf1, with associated *r*^2^ values of 0.98 and 0.94 (Tab. S4, Fig. 3*A*, 3*B*). These improvements, corresponding to ∼5% and 24% increases in explanatory power over respective mononucleotide models, are consistent with the potential for physically interacting nucleotides to affect binding energies through local structural distortions. Considering all possible dinucleotide features accounts for nearly all of the remaining variance in NN-derived binding energies (improvements of approximately 1% and 5% for Pho4 and Cbf1, respectively). These findings highlight the differential degree to which epistasis defines binding even amongst structurally related TFs, which ultimately determines the predictive power and accuracy of widely-used PWMs.

To visualize and interpret binding energy contributions of dinucleotides alone, we calculated the mean residual ΔΔG from the linear regression against PWM-predicted ΔΔG values for all possible dinucleotides both within and across flanking sequences (Fig. 4*C*). Nucleotide interactions that exhibit measured ΔΔGs lower than expected based on considering the linear combination of individual mononucleotides are positively epistatic; conversely, interactions that exhibit negative epistasis increase measured ΔΔGs more than expected. The largest magnitude epistasis is observed for dinucleotides immediately upstream or downstream of the E-box (N4/N5 or N6/N7 pairs), with absolute energetic differences among combinations spanning ∼0.5 kcal/mol (approximately 1 RT), and epistatic interactions primarily occur within flanks rather than between them. Both inter- and intra-flank dinucleotides exhibited palindromic arrangements near the core motif, consistent with the expectation of binding site symmetry in homodimeric TFs like Pho4 and Cbf1 (61, 71). For Cbf1, TT and TG dinucleotides upstream of the motif (and the corresponding downstream panlindromes) exhibit large magnitude positive and negative epistasis, respectively. Although the overall magnitude of epistasis is significantly smaller for Pho4, a GG dinucleotide downstream of CACGTG shows strong synergistic effects. Interestingly, the Pho4 crystal structure reveals direct contacts between the Arg2 and His5 residues and this same GG dinucleotide, providing a potential structural basis for this observation (61).

**Fig. 4.**
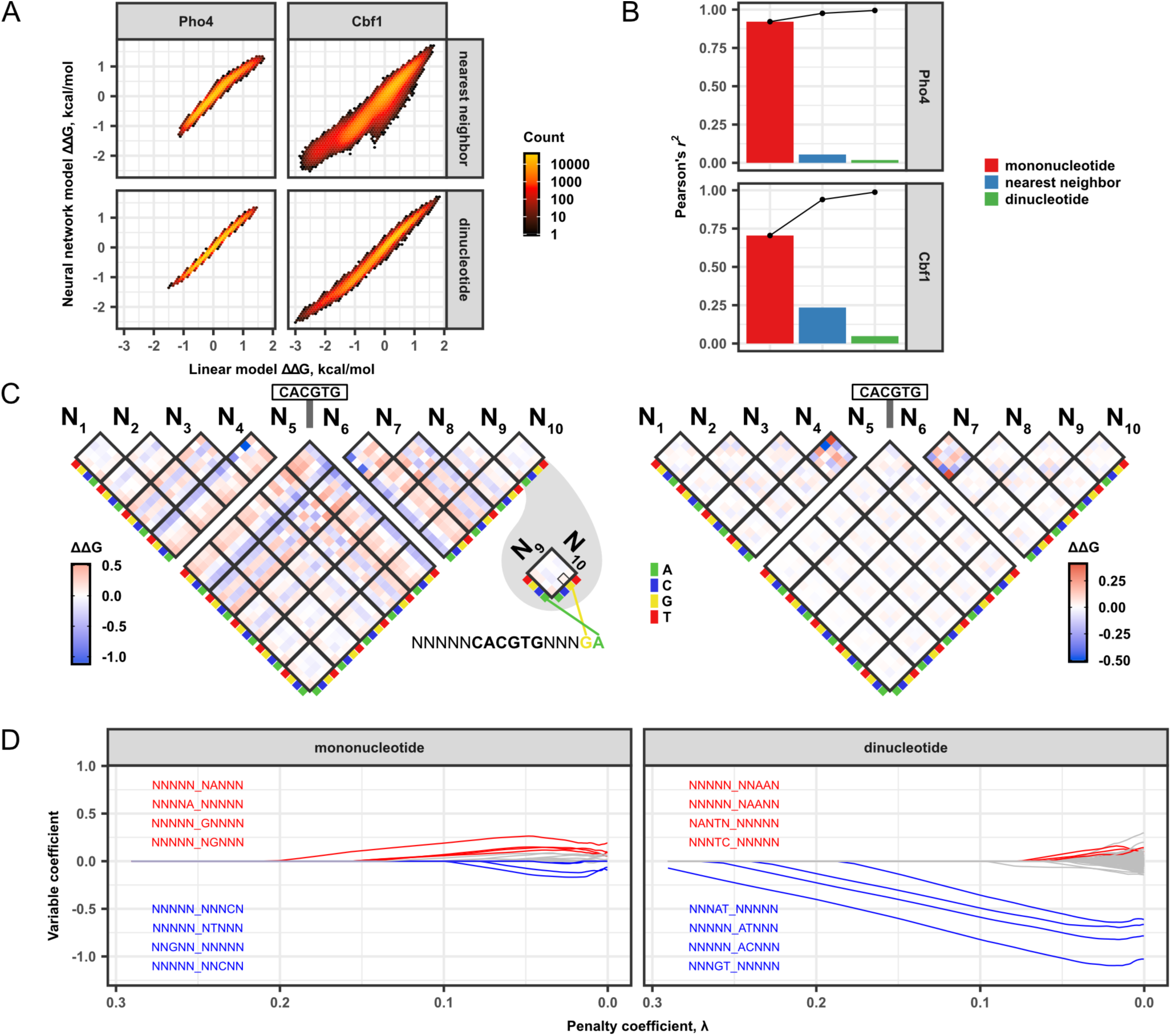
NN model interpretation using dinucleotide features. (*A*) For Pho4 and Cbf1, density scatter plots show the correlation between NN model estimates and dinucleotide (nearest neighbor = only adjacent pairs) additive model predictions based on those estimates. (*B*) Variance in Pho4 and Cbf1 NN model estimates, captured by squared Pearson’s *r*^2^ compared to linear models predictions, attributed to sequence feature groups. (*C*) Cbf1 heatmap of mean energetic contribution for all dinucleotide configurations (left) and that of mean residual energetic contributions when mononucleotide effects are removed (right). Color scale: blue = high affinity, red = low affinity. (*D*) Cbf1 LASSO regression penalization traces arranged by type of model feature. Lines represent magnitude of individual sequence feature coefficients as a function of penalty coefficient λ. Labels show flanking sequence feature. Color scale: blue = four most persistent negative energetic contributions, red = four most persistent positive energetic contributions.

### Incorporating weight constraints into dinucleotide models confirms that Cbf1 interactions are significantly more epistatic

Models that incorporate additional free parameters should always increase explanatory power. While mononucleotide models attempt to describe all 1,048,576 observed measurements using only 40 free parameters (4 nucleotides per position across 10 positions), the nearest-neighbor dinucleotide model adds another 128 free parameters (16 pairs across 8 positions), and the all dinucleotides model includes 720 free parameters representing the contribution from all combinations of nucleotide identities [4^2^ =16] and positional pairs 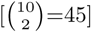. In most cases, dinucleotide coefficients are near zero (Fig. 4*C*), meaning that they contribute little explanatory power. To systematically identify a minimal set of important features that define sequence specificity in an unbiased fashion, we used least absolute shrinkage and selection operator (LASSO) regression to develop parsimonious linear models with weight constraints (72). Briefly, non-zero coefficients in the model are penalized, leading to inclusion of only the most explanatory variables with respect to reduction in squared error (Fig. S11). The regression explores a range of penalization stringencies to distinguish important sequence features based on differential coefficient minimization rates.

We applied LASSO regression to systematically select a parsimonious set of sequence features with optimal explanatory power among all possible dinucleotide features. From the selected Cbf1 features, four nearest neighbor dinucleotides exhibited relatively large initial coefficient magnitudes and persisted as model variables throughout most of the penalization regime. These four sequence features group into two pairs of palindromic dinucleotides spanning the core motif: (NNNAT_NNNNN, NNNNN_ATNNN) and (NNNGT_NNNNN, NNNNN_ACNNN) (Fig. 4*D*). Strikingly, this set of dinucleotide features exhibits model coefficients that are up to 3-fold greater in magnitude than that of the most significant mononucleotide feature (Fig. 4*D*, S12), highlighting the importance of dinucleotides to Cbf1 binding specificity in particular. Among the selected dinucleotide features for both Pho4 and Cbf1, nearest neighbor pairs exhibited the largest coefficient magnitudes (Fig. S12). Short-range nucleotide interactions contribute most to higher-order substrate specificity, consistent with the feature preferences of yeast TFs determined by ChIP-chip (45).

### Orthogonal *in vitro* biochemical measurements confirm results obtained via high-throughput sequencing

To confirm that NN model predictions provide accurate per-sequence estimates of true binding energies, we quantitatively compared titration-based ΔΔG values with unprocessed measurements and NN predictions. Using traditional fluorometric MITOMI, we determined the ΔΔGs for Pho4 and Cbf1 binding to singlesite variants of the ACAGA_TCGAG flanking sequence (Tab. S3, Fig. S6*A*, S6*B*). In addition, we correlated NN predictions with a set of previously reported ΔΔG measurements of CACGTG flanking site mutations (17, 31). Consistent with the Monte Carlo simulations, ΔΔG values calculated directly from raw sequencing data showed essentially no correlation to direct measurements, with *r*^2^ values ranging between 0.07–0.16 and 0–0.24 for Pho4 and Cbf1, respectively (Fig. S13). NN-predicted values showed remarkable agreement, with *r*^2^ values ranging between 0.76–0.94 and 0.61–0.69 for Pho4 and Cbf1. In all cases, predictions agreed with observations within ∼1 kcal/mol. Taken together, these results establish that the Pho4 and Cbf1 NN models presented here yield a complete and accurate measurements of binding energies for >1 million TF-DNA interactions with similar resolution to ‘gold-standard’ biochemical measurements.

### High-resolution *in vitro* affinity measurements can be used to identify biophysical mechanisms underlying *in vivo* behavior

The role of transcriptional activators *in vivo* is not simply to bind DNA, but rather to bind specific genomic loci and regulate transcription of downstream target genes. The high resolution of these comprehensive binding energy measurements makes it possible to quantitatively estimate the degree to which measured binding affinities explain differences in measured TF occupancies, rates of downstream transcription, and ultimate levels of induction.

First, we compared NN-modeled ΔG values with measured rates of transcription and fold change induction for engineered promoters containing CACGTG Pho4 consensus sites with different flanking sequences driving the expression of fluorescent reporter genes (16, 17). As reported previously, rates of transcription and induction scaled with measured ΔG values that spans −1.5 kcal/mol (Fig. S14). Next, we compared NN-derived binding energies with reported levels of TF occupancy *in vivo* at CACGTG consensus sites in the *S. cerevisiae* genome for both Pho4 and Cbf1 (62). Large magnitude TF enrichment induced by phosphate starvation was observed at loci with measured K_d_ values of around 100 nM or lower (Fig. 5*A*). While TF enrichment roughly correlated with binding energy, very high affinity sequence showed strikingly low enrichment. The observed nonlinearities may indicate the degree to which other regulatory mechanisms, such as cooperation and competition among TFs or changes in DNA accessibility due to nucleosome positioning, contribute to reported TF enrichment (62). Alternatively, these nonlinearities may reveal the need for higher resolution *in vivo* measurements to test the degree to which binding energies alone dictate occupancies.

**Fig. 5.**
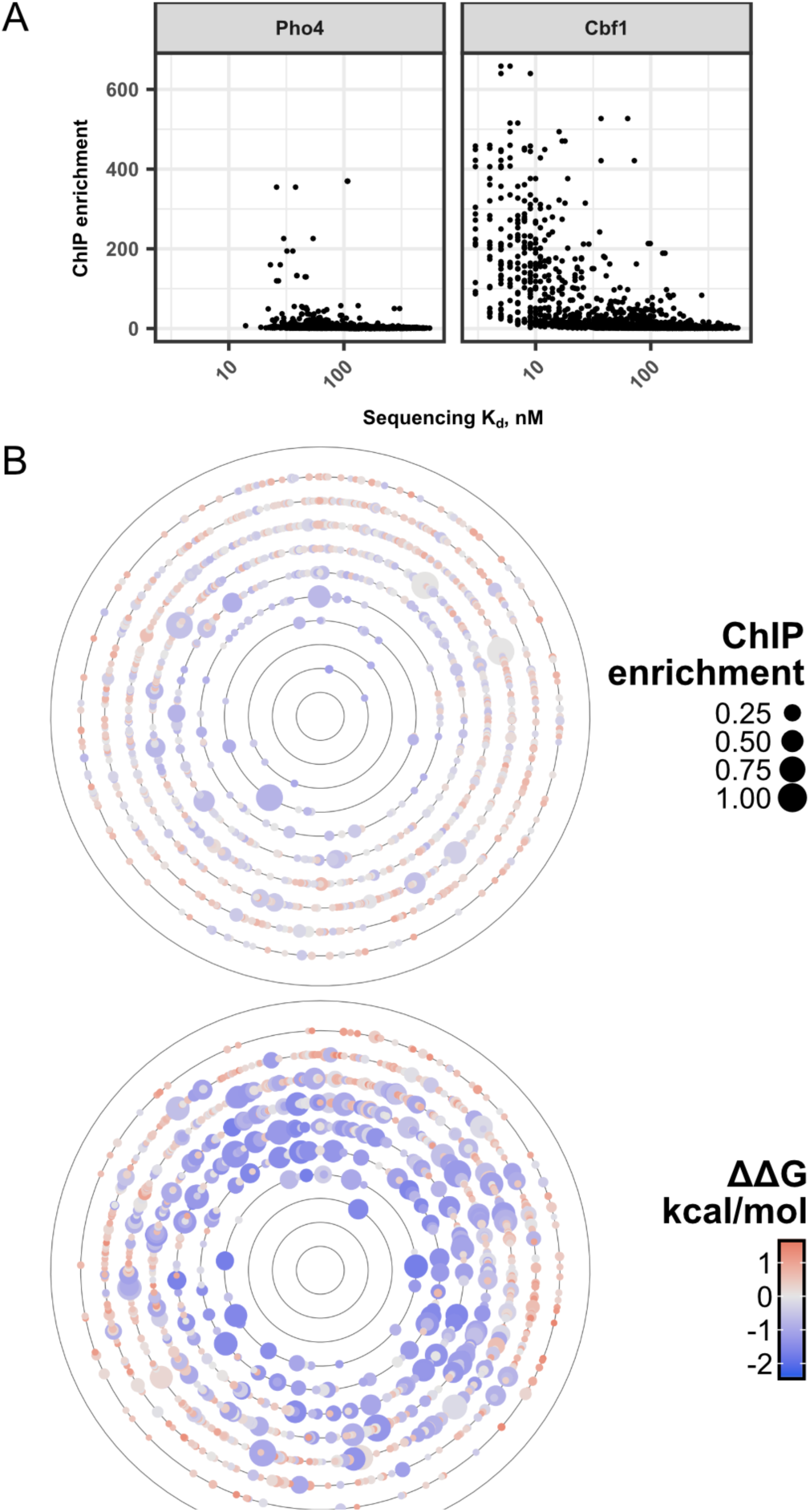
Large-scale Pho4 and Cbf1 binding energies and *in vivo* activity. (*A*) Pho4 and Cbf1 equilibrium binding constants (K_d_, in nM) compared to *in vivo* enrichment determined by ChIP-seq (62). (*B*) Functional-energetic landscapes of differential relative ChIP enrichment at dynamically regulated loci, relative to the measured highest affinity sequence in Hamming distance space. Point size: large = high enrichment, small = low enrichment. Color scale: blue = high affinity, red = low affinity.

To better understand the relationship between binding energies and *in vivo* occupancies, we visualized the complete binding energy landscapes for both Pho4 and Cbf1 as a function of sequence space (Fig. 5*B*, S15). The single highest affinity sequence for each TF was placed at the center of a series of concentric rings, each of which includes all sequences at a given Hamming distance from the highest affinity sequence. Within each ring, points representing each sequence are arranged in alphabetical order, with the color of each point reporting the measured ΔΔG for that sequence. As expected, the energetic landscape forms a somewhat rugged funnel, with binding energies increasing monotonically on average with mutational distance from the highest-affinity site (Fig. S16). Next, we projected flanking site occupancies from ChIP-seq experiments (62) onto these binding affinity landscapes to yield a composite functional-energetic landscape (Fig. 5*B*). For both Pho4 and Cbf1 the majority of enriched genomic loci are greater than four mutational steps away from the the global minimum, corresponding to mean increases in binding energy of approximately 0.8 and 1.5 kcal/mol, respectively (Fig. S17). These quantitative comparisons between measured affinities and *in vivo* occupancies establish that even relatively small differences in ΔΔG are associated with differential TF enrichment.

## Discussion

TFs play a central role in regulating levels of gene expression throughout development and allowing organisms to adapt to changing environmental conditions. The ability to quantitatively predict levels of TF occupancy *in vivo* from DNA sequence would therefore be transformative for our understanding of cellular function. Given that the probability of TF occupancy at a given locus includes an exponential dependence on the corresponding TF-DNA binding energy (73), accurate occupancy predictions require the ability to resolve even small (∼1-2 kcal/mol) changes in binding energies at high resolution. Towards this goal, we developed a novel assay designed to provide comprehensive and quantitative measurements of near-neutral changes in binding energies caused by mutations in the flanking sequences surrounding TF consensus target sites. By training a neural network (NN) on estimates of binding energies for millions of sequences, we obtained a model that incorporates all higher-order complex interactions required for accurate binding energy estimates for each sequence.

We find that a wide variety of mutations in flanking nucleotides outside of “core motifs” (as defined by consensus PWMs) can change binding energies by an amount equal to or greater than mutations within the core consensus. For example, the difference in binding energy between a *TCCCC* **CACGTG***CCCCA* sequence and a *AATTT* **CACGTG***AAAAG* sequence is ∼2.6 kcal/mol, equivalent to mutating the central consensus site from **CACGTG** to **C <u>GT</u> GTG**. However, current representations of TFBSs would predict a change in binding energy for only the core mutation. This discrepancy has several implications for our understanding of transcriptional regulation *in vivo*. First, flanking sequence effects may explain mysteries regarding chromatin immunoprecipitation (ChIP) data in which some genomic loci are occupied despite an apparent lack of a consensus site while other accessible regions containing consensus sites remain unoccupied. Second, many current efforts to infer the presence of bound TFs first analyze DNAse-seq or ATAC-seq data to identify regions of accessible DNA, and then scan these regions for putative bound TFs by searching for sequence similarities to known TFBSs. Failing to consider the effects of flanking sequences could return a significant number of both false positives (in which a consensus site with unfavorable flanking nucleotides is considered to be bound) and false negatives (in which a mutated consensus site has strongly favorable flanks but is considered unoccupied).

In practice, measuring complete binding energy landscapes remains rare, with most assay development focused towards discovery of the highest-affinity sequences. The quantitative and complete binding energy landscapes presented here provide a unique opportunity to explore the mechanisms that drive evolution of transcriptional regulatory networks. It has been hypothesized that high affinity, but sub-maximal, TF binding sites may be evolutionarily favorable due to the potential for greater dynamic transcriptional control (74). Consistent with this hypothesis, we find that the most highly occupied sites *in vivo* are mutationally distant from the highest affinity flanking sequences. These large mutational distances could indicate an evolutionary buffer employed by an organism to avoid sequence proximity to a sub-optimal binding extreme. In addition, elevated levels of epistasis are thought to produce more rugged energetic landscapes compared to those created by additive binding interactions (75). Given that epistatic dinucleotide interactions play a larger role in determining Cbf1 binding specificity, we speculate that Cbf1 binding sites can traverse fewer non-deleterious evolutionary pathways than Pho4, ultimately rendering Pho4 binding sites more evolutionarily plastic than those of Cbf1.

Systematic comparisons between per-sequence estimates of binding energies output by a NN and those output by a series of linear models revealed the mechanistic features that drive specificity and quantified their contributions to observed binding energies. These results have relevance to recent debates surrounding the relative utility of DNA sequence-based models (PWMs) and DNA shape-based models representing TF specificity. Both models parameterize DNA binding preferences by a set of four values at each position: PWMs represent specificities based on preferences for each of the DNA bases (A, C, G, and T) (35), while shape-based models express preferences in terms of minor groove width, propeller twist, helical twist, and roll (50–54). While these models significantly enhance the ability to extract mechanistic determinants of specificity from sparse data, higher-order information is lost in the process. Here, we demonstrate that models based on nearest-neighbor dinucleotide preferences are sufficient to fully explain observed binding behavior, consistent with biophysical observations that local DNA structure is largely determined by base stacking interactions and inter-base pair hydrogen bonds in the major groove between adjacent base pairs (54, 76, 77). Such nearest-neighbor dinucleotide models require only a modest increase in the number of required free parameters relative to mononucleotide models. While the NN’s capacity to incorporate higher-order complexity ultimately proved unnecessary for accurately modeling Pho4 and Cbf1 binding specificities, high-resolution predictions output by the NN were essential to quantify the degree to which simpler models could explain observed behavior. The high resolution of these measurements further allows direct quantification of the degree to which thermodynamic models based on the binding energies can and cannot predict behavior *in vivo*.

The simulation-guided assay design and experimental assay presented here should allow a much broader diversity of labs to make comprehensive and high-resolution measurements of binding energy landscapes. While the assay is deployed here for a specific use case (high-resolution measurements of near-neutral effects over a small energy range), these simulations can guide choice of sequencing depths to resolve absolute binding energies across a variety of applications and platforms (9, 11, 28, 78, 79), including target site discovery efforts. The experimental assay itself further offers the resolution of traditional MITOMI or HiTS-FLIP fluorescence-based assays while requiring significantly less equipment and infrastructure. Traditional MITOMI fluorescence assays require both a DNA microarray printer to deposit DNA on slide surfaces and either a high-cost fluorescence scanner or fully automated microscope capable of quantitative fluorescence imaging of a slide with a microfluidic device attached; HiTS-FLIP assays require access to a customized Illumina GAIIx sequencing platform. The use of a sequencing-based readout eliminates these requirements, allowing any laboratory with access to educational or commercial deep sequencing services to measure energies at this scale and resolution. Moreover, the valving required on the microfluidic device itself is significantly simpler than for traditional MITOMI assays, reducing the pneumatics infrastructure required to operate devices.

Finally, this assay provides unique opportunities in future work to probe additional control mechanisms that influence TF binding *in vivo*. Introduction of synthesized DNA libraries containing modified bases involved in epigenetic regulation (*e.g.* 5-methylcytosine, 5-hydroxymethylcytosine, 5-formylcytosine, 5-carboxylcytosine) could allow systematic investigation of how these modifications affect TF specificities. The assay should further be compatible with DNA libraries assembled into nucleosomal arrays *in vitro*, facilitating direct and quantitative investigation of how competition between TFs and nucleosomes dictates occupancies, the degree to which particular TFs act as “pioneer factors”, and how site-specific histone modifications (*e.g.* methylation, acetylation, phosphorylation, ubiquitinylation) (80–82) influence this competition. It may also be possible to measure relative binding energies of TF interactions with extracted DNA or chromatin derived from cells under defined stimulation (*e.g.* drug response, disease) (83). Taken together, these areas of study highlight the complex interplay of chromatin landscape and TF binding, aspects of which may be characterized in the future using a variant of the assay and analysis pipeline presented in this work. In addition, the simulations presented here provide guidelines for the development of sequencing-based assays designed to measure binding energies for a wide variety of macromolecular interactions, including both protein-RNA and protein-protein interactions.

## Materials and Methods

### Binding energy calculations

Using the model proposed by Djordjevic, *et al.* (56) that was refined by Stormo and colleagues (10, 39), we consider the binding of DNA sequences *S*_*i*_ among many *S*_*j*_ competitor substrates at equilibrium with a transcription factor. The probability of being bound to the transcription factor as a function of sequence identity, represented by *P* (*bound | S*_*n*_), is given by:

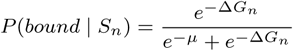

Where Δ*G*_*n*_ is the binding free energy and *μ* is the chemical potential, equal to the natural log of the transcription factor concentration. Both terms are in units of RT.

Sequencing of TF-bound substrates yields *P* (*S*_*i*_ *| bound*), which is the inclusion probability of *S*_*i*_ among the bound substrate distribution. From the partition probability of substrates between bound and unbound, *P* (*S*_*i*_ *| bound*), application of Bayes’ theorem and the law of total probability yields:

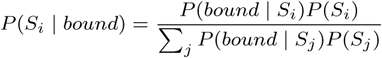

Where *P* (*Sn*) describes the probability of a given species within the input substrate distribution. The combination of these two equations returns *P* (*S*_*i*_ *| bound*) as a function of binding energies and input probabilities.

Next, we make the following assumptions:

- The transcription factor concentration is substantially lower than the total DNA concentration.
- As the TF concentration is minimized, *μ* approaches negative infinity. In effect, this causes the denominator in equation 1 to be dominated by the *e*^*-μ*^ term, which we can approximate with a constant *C ≃ e*^*−μ*^ + *e*^*ΔG*^.
- The sum of *P* (*S*_*j*_) in a large population is equal to one.

Applying these assumptions to equation 2, it becomes possible to isolate Δ*G*_*i*_ as a function of *P* (*S*_*i*_ *| bound*) and *P* (*S*_*i*_). This equality is given in equation 3, in units of RT:

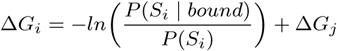

Where Δ*G*_*j*_ equals the total binding energy excluding contribution from *S*_i_. Setting this value to zero yields the relative binding affinity of *S*_*i*_, represented by ΔΔ*G*_*i*_ in units of RT, which is given in equation 4:

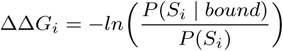

### Neural network binding models

The input for the neural network was defined as a flattened 40-element vector derived from a 4x10 onehot encoded matrix. Within the initial matrix, each row represents a nucleotide species (A, C, G, or T) and each column represents a position within the ten nucleotide sequence flanking the E-box motif. The output of the neural network was defined as a scalar corresponding to the predicted ΔΔG value for the species of interest. The network consisted of three hidden layers of size 500, 500, and 250 units, respectively. All weights were initialized with Xavier initialization (84) and all layers used batch normalization (85) and ReLU activation. The entire available sequence space was randomly divided into training (60%), validation (10%), and test (30%) datasets. The networks were trained on the training portion of the data using stochastic gradient descent until the validation set root mean squared error failed to decrease for three consecutive epochs. At this point, the learning rate (initialized at 10*≠*3) was decreased in 10-fold increments, and training continued until error failed to improve for a further two epochs when training was halted to prevent overfitting (Fig. SXX).

### Linear binding models

Energetic models of TF substrate specificity use linear combinations of sequence features to explain variation in binding energy (70). All linear binding models presented in this work were training on the binding predictions Our mononucleotide model includes sequence features consisting of DNA bases present at each flanking position. We parameterized the nearest neighbor dinucleotide models using all mononucleotide features plus all possible combinations of adjacent nucleotide pairs. The full dinucleotide model builds upon the nearest neighbor model with the addition of all non-adjacent (gapped) dinucleotide combinations. All linear binding models were trained using the same 60% of the sequence space as for the neural network models. Reported accuracies are calculated with respect to the held-out 40% of the sequence space.

## ACKNOWLEDGMENTS

We thank Justin Kinney for library design discussions, Hua Tang for discussions of epistasis and statistics, Anshul Kundaje for neural network discussions, and Daniel Herschlag and Rhiju Das for helpful comments on the manuscript.T.C.S acknowledges support from an NSF Graduate Research Fellowship. This work was supported by NIH/NIGMS grant R00GM09984804. P.M.F. also acknowledges support from the Sloan Research Foundation, and a McCormick and Gabilan faculty fellowship. P.M.F. is a Chan Zuckerberg Biohub Investigator.

